# Relationship between MEG global dynamic functional network connectivity measures and symptoms in schizophrenia

**DOI:** 10.1101/432435

**Authors:** L. Sanfratello, J.M. Houck, V.D. Calhoun

**Author notes:** **Corresponding Author:** Lori Sanfratello, 1101 Yale Blve NE, Albuquerque, NM 87059 USA.

## Abstract

An investigation of differences in dynamic functional network connectivity (dFNC) of healthy controls (HC) versus that of schizophrenia patients (SP) was completed, using eyes-open resting state MEG data. The MEG analysis utilized a source-space activity estimate (MNE/dSPM) whose result was the input to a group spatial independent component analysis (ICA), on which the networks of our MEG dFNC analysis were based. We have previously reported that our MEG dFNC revealed that SP change between cognitive meta-states (repeating patterns of network correlations which are allowed to overlap in time) significantly more often and to states which are more different, relative to HC. Here, we extend our previous work to investigate the relationship between symptomology in SP and four meta-state metrics. We found a significant correlation between positive symptoms and the two meta-state statistics which showed significant differences between HC and SP. These two statistics quantified 1) how often individuals change state and 2) the total distance traveled within the state-space. We additionally found that a clustering of the meta-state metrics divides SP into groups which vary in symptomology. These results indicate specific relationships between symptomology and brain function for SP.

## 1. Introduction

Numerous psychiatric disorders, including schizophrenia, have low treatment success rates and create a diminished quality of life for those that suffer. Schizophrenia is a debilitating mental disorder whose treatment has changed little in decades. Successful treatment is often focused on symptom reduction yet many individuals continue to have deeply unsatisfying dayto-day functioning. It is imperative therefore that research capitalize upon newer techniques and analysis methods that can aid in informing: identification and classification, such as patients with co-morbidities; new treatment targets; and illness trajectory. We show here how dynamic functional network connectivity (dFNC) utilizing MEG data may provide unique information that moves us toward a better understanding of the relationship between cognitive states, how individuals move through these states, and symptoms in SP.

Functional network connectivity (FNC) is defined as the way in which sets of brain areas (or networks) work together over time, represented by statistical associations between the networks where the spatial proximity of the regions to one another is unimportant. Since an increasing body of literature suggests that neural oscillations perform a key role in binding separate brain regions together and promoting information transfer between distant brain areas^1–3^ FNC has become an important metric for the study of how this process naturally occurs^4–6^. Furthermore, appropriate connectivity between brain regions is now generally accepted as being key to healthy brain function^7^. Clearly the temporal as well as the spatial properties of these networks is important to our understanding of brain function, and it is probable that the definition of a network may vary on different time scales^8^. FNC has often been investigated within the resting state (i.e., in the absence of a defined task) using fMRI^9–11^ and, to a lesser extent the electrophysiological methods MEG and EEG^12–15^, in diverse populations including schizophrenia, depression, bipolar disorder, and in aging^16–25^. However, the timescale of an FNC map (the scan length) is arbitrary, and there is evidence that important information is being “diluted” with the use of FNC maps that condense several minutes of measurement into a single statistical map. This is particularly a problem for electrophysiological data where there is evidence of brain-state changes on the order of hundreds of milliseconds^26^.

A logical extension of FNC that looks at how states vary over small time “windows” in order to capture networks on a finer temporal scale, has been termed dynamic functional network connectivity (dFNC). dFNC is defined as the (Gaussian tapered) *windowed* zero-lag cross-correlations among networks. Much previous work has found that these sets of correlations can be grouped into stable repeating patterns of activity representing a few often visited brain states that may be identified using a k-means clustering of the correlation patterns at each time-point (cluster states^11,27^). dFNC has also been investigated in resting state fMRI^27,28^ and in diagnostic groups such as schizophrenia patients^20,27,29^. The statistics (meta-state statistics) derived from these repeating connectivity patterns can then be used to further investigate population differences. We here determined the relationship between meta-state statistics and symptomology in SP. We hypothesized that the meta-state statistics where we found significant differences between HC and SP groups in our previous study^34^ would correlate with symptoms for SP. These metrics measured how often SP changed between brain states and how different the next brain state was from the previous one occupied (**Table 1**). We also completed a preliminary investigation of whether global-level statistics would group SP in a data-driven fashion, where the number of groups was not specified a priori.

**Table 1.** HC (N = 36) and SP (N = 36) groups reveal significant differences in how often they change between brain states (“Change between states”) and in how different the previous brain state is from the next one occupied (“Total distance traveled through state space”). Significant differences between HC and SP groups are highlighted (p < 0.05).

|  | Number of states | Change between states | State space span | Total distance traveled through state space |
| --- | --- | --- | --- | --- |
| mean HC | 10.11 | 50.39 | 5.25 | 51.53 |
| mean SP | 11.03 | 58.89 | 5.56 | 61.05 |
| T-value | -1.42 | -2.15 | -1.43 | -2.33 |
| p-value | 0.16 | **0.035\*** | 0.16 | **0.022\*** |

## 2. Methods

### 2.1 Participants

Briefly, this investigation utilized existing data^30^ from 36 schizophrenia patients from whom informed consent was obtained according to institutional guidelines at the University of New Mexico Human Research Protections Office (HRPO). Note that these were the same individuals used for the analysis presented in **Table 1**^34^. Note that there was no significant difference in age or gender between the HC (Mean Age+/−SD = 35.7+/−11.8; 27M/9F) and SP (Mean Age+/−SD = 38.4+/−13.5; 31M/5F) groups and all participants were right handed. All participants were compensated for their participation. Patients with a diagnosis of schizophrenia or schizoaffective disorder were invited to participate. Each patient completed the Structured Clinical Interview for DSM-IV Axis I Disorders^31^ for diagnostic confirmation and evaluation of co-morbidities. Exclusion criteria included history of neurological disorders, mental retardation, substance abuse, or clinical instability. Patients were treated with a variety of antipsychotic medications, therefore doses of antipsychotic medications were converted to olanzapine equivalents^32^. Each participant completed resting MEG and structural MRI scans; some also completed additional task and resting fMRI scans as part of a larger study. All patients also completed the positive and negative syndrome scale^33^ (PANSS). Positive symptoms: delusions, conceptual disorganization, hallucinations, excitement, grandiosity, suspiciousness/persecution, hostility; Negative symptoms: blunted affect, emotional withdrawal, poor rapport, passivity, difficulty in abstract thinking, lack of spontaneity and flow of conversation, stereotyped thinking. Mean PANSS scores for SP were (Total = 30.3+/−6.6; Positive = 15.2+/−5.0; Negative = 15.1+/−5.6). The participant data and preprocessing used for this study overlapped with that presented in [17, 34] but the analytic approach developed in this work and the questions being investigated are novel and distinct.

### 2.2 MEG

The details of the MEG data acquisition and processing can be found in [34] as well as in the supplementary material. Briefly, five minutes of eyes-open resting state MEG data were acquired. The cortical surface of each participant was reconstructed from T1-weighted MRI images. Activity at each vertex of the cortical surface was determined using a noise-normalized minimum norm estimate (MNE/dSPM)^35^. In essence, MNE/dSPM identifies where the estimated current differs significantly from baseline noise (e.g., empty room data); this method also acts to reduce the location bias of the estimates^36^. Spatiotemporal source distribution maps downsampled to a 50 Hz sampling rate were obtained at each time point (providing an upper frequency bound of 25 Hz) for 60 sec of the data. Group spatial ICA (gsICA) was applied to the individual subject MNE/dSPM source-space maps using the GIFT toolbox (http://mialab.mrn.org/software/gift) as in our prior work^17,34^ in order to mitigate the signal leakage issue (i.e. the leakage of signal between projected timecourses, or point spread, which manifests as zero-lag correlations between timecourses of spatially separate regions). This returned to us 32 independent components (ICs or “networks”), which were determined to be non-artifactual.

### 2.3 dFNC and k-means clustering of repeating cognitive states

By definition a dFNC is the (Gaussian tapered) *windowed* zero-lag cross-correlations among brain networks. Due to the frequency content of our data we chose a 4sec window for the dFNC, capturing all available frequencies of interest (1Hz-25Hz). Next, a dFNC analysis was conducted between all 32 ICs calculated from the gsICA at each timepoint. The resulting states were then clustered into repeating patterns using the k-means clustering method^11,27^. This resulted in 3 cluster-states whose information was then summarized by allowing the states to overlap in time to create meta-states. The meta-state metrics calculated in this manner were^27^: 1) The number of distinct meta-states subjects occupy during the scan length (“Number of states”); 2) The number of times that subjects switch from one meta-state to another (“Change between states”); 3) The range of meta-states subjects occupy, i.e., the largest L1 distance between occupied meta-states (“State span”); and 4) The overall distance traveled by each subject through the state space (the sum of the L1 distances between successive meta-states, (“Total distance”). These meta-state statistics were then investigated for their relationship to symptoms in SP.

### 2.4 Investigation into the relationship between meta-state statistics and symptomology in SP

Since we were interested in determining how symptoms are related to how SP traverse cognitive states we correlated total positive and total negative symptoms with each of the 4 meta-state statistics. In addition, one goal of our work is to develop sensitive tools to classify individuals into appropriate groupings, particularly when individuals have co-morbidities or symptomology that overlap with multiple diagnoses. Therefore, we also investigated if a data-driven k-means clustering of the meta-state level statistics would inform classification of patients.

## 3. Results

### 3.1 dFNC meta-state statistics and symptoms in SP

We found that positive symptoms significantly correlate with the meta-state statistics “Change between states” and “Total distance” (**Fig. 1**). No significant correlation between negative symptoms and the various meta-state statistics was found. Results from a data-driven k-means cluster analysis based on the global meta-state level statistics revealed a trend for patients to group according to their symptomology. Specifically, the analysis determined that 4 groups was optimal for this dataset (> 80% variance explained, **Table 2**), where it revealed one group that was high in reported positive and negative symptoms (group 2), one group which was relatively high in positive symptoms but low in negative symptoms (group 4), one which was relatively low in positive symptoms but high in negative symptoms (group 1), and one group which was relatively low in both positive and negative symptoms (group 3). We find significant differences in negative symptoms between groups 1 & 4 and 2 & 4, as would be expected if this is to be a useful method for differentiating between patient symptomology. However, we do not find significant differences between any of the groups for positive symptoms for this small set of patients. It is also clear from **Table 2** that the 4 groups are significantly different from each other on the meta-state statistics “Change between states” and “Total distance”, and that in addition groups 1 & 2 are significantly different from each other on the metric “Number of states” and that groups 2 & 3 and groups 3 & 4 are significantly different from each other on the meta-state metric “State span”. No significant differences were found between the 4 groups in age, gender, or medication dose.

**Table 2.** Clustering of meta-state statistics splits SP into groupings differentiated by symptomology. Significant differences between groups are highlighted (p < 0.05).

|  | # SP | Gender<br>0=f; 1=m | Age | Olanzapine<br>Equivalent | total<br>positive | total<br>negative | Number of<br>states | Change bw<br>states | State span | Total<br>distance |
| --- | --- | --- | --- | --- | --- | --- | --- | --- | --- | --- |
| Group 1 | 9 | 0.89 | 33.19 | 19.82 | 13.00 | 18.78 | 9.89 | 35.56 | 5.22 | 37.11 |
| Group 2 | 7 | 0.86 | 40.38 | 14.53 | 16.86 | 18.29 | 12.14 | 83.43 | 6.00 | 88.00 |
| Group 3 | 7 | 0.86 | 44.29 | 12.11 | 14.57 | 13.86 | 10.00 | 50.86 | 5.00 | 51.29 |
| Group 4 | 13 | 0.92 | 37.85 | 15.05 | 16.15 | 11.46 | 11.77 | 66.00 | 5.85 | 68.38 |
| ttest 1-2 |  | 0.8610 | 0.3720 | 0.3695 | 0.2081 | 0.8628 | **0.0449** | **6.59E-010** | 0.0828 | **1.64E-010** |
| ttest 1-3 |  | 0.8610 | 0.0654 | 0.1666 | 0.3773 | 0.0900 | 0.9273 | **2.67E-004** | 0.6821 | **3.56E-004** |
| ttest 1-4 |  | 0.7960 | 0.4363 | 0.2698 | 0.1272 | **0.0007** | 0.1109 | **1.25E-010** | 0.0694 | **2.12E-010** |
| ttest 2-3 |  | 1.0000 | 0.6027 | 0.6122 | 0.4702 | 0.1616 | 0.0554 | **2.60E-009** | **0.0213** | **4.87E-010** |
| ttest 2-4 |  | 0.6601 | 0.7282 | 0.8970 | 0.8055 | **0.0030** | 0.7381 | **4.69E-008** | 0.2986 | **9.08E-008** |
| ttest 3-4 |  | 0.6601 | 0.2642 | 0.4116 | 0.4484 | 0.2223 | 0.1616 | **1.70E-007** | **0.0129** | **3.97E-007** |

**Fig. 1.**
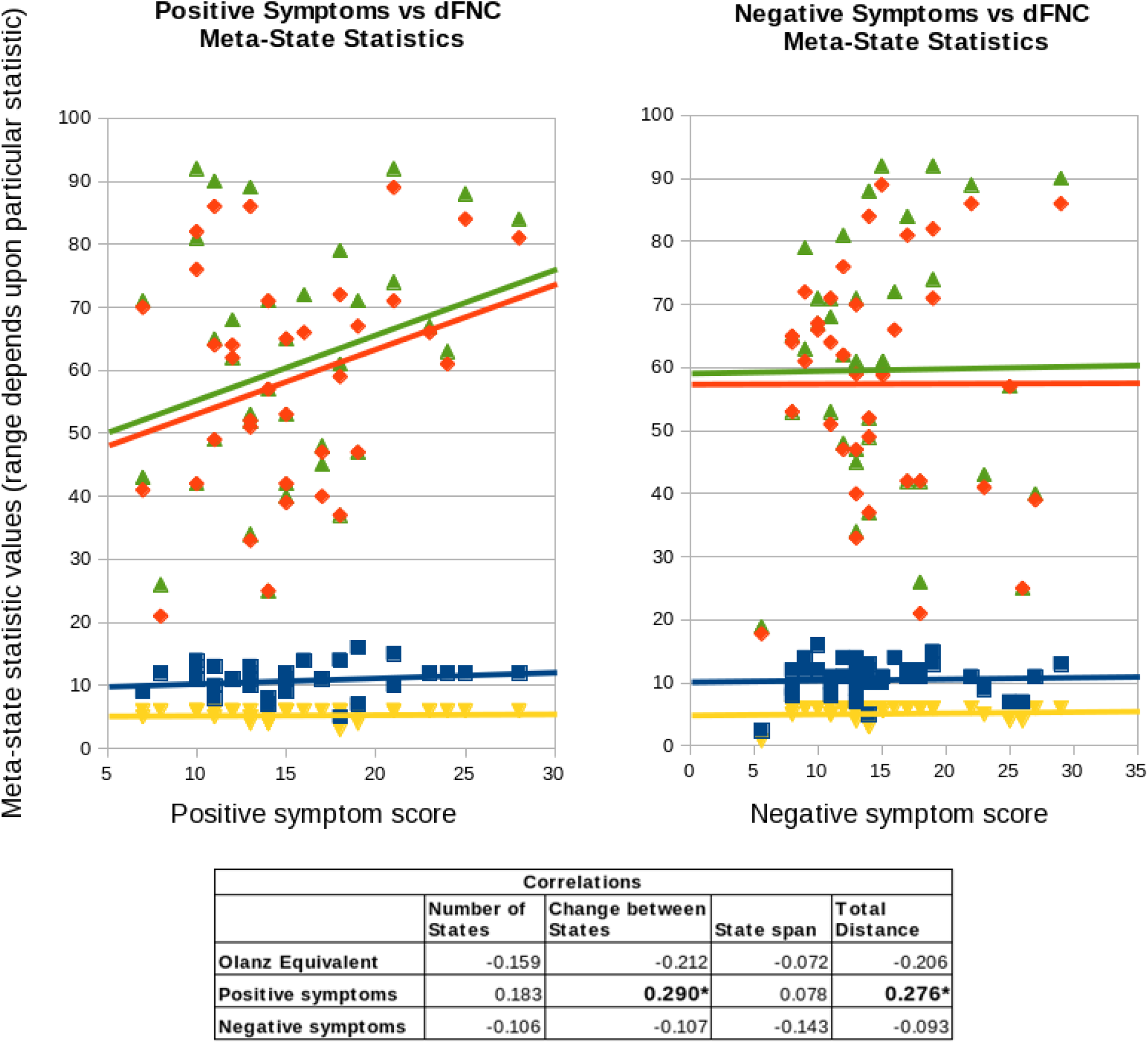
Correlations between global meta-state statistics and symptoms in SP. Blue squares and line = Number of states; Orange diamonds and line = Change between states; yellow down pointing triangles and line = State span; green up pointing triangles and line = Total Distance. Significant correlations (p<0.05) are observed for positive symptoms and meta-state statistics “change between states” and “total distance traveled”.

## 4. Discussion

Perhaps the most important message from the current results is that for SP positive symptoms are significantly correlated with how patients are transitioning between brain states. Specifically, the more positive symptoms a patient has the *more likely they will be to both change states frequently and to change to states which are more different from the previous state occupied*. This has implications for treatment, especially if these results can be replicated in a larger sample and with patients who have a broader range of symptom scores (scores in the current cohort range from 7 to 28 for positive symptoms and from 7 to 29 for negative symptoms, where only 5 individuals showed scores > 20). The particular relationship we observed, where positive symptoms are positively correlated with how often individuals change states, may have a direct relationship with the well-documented difficulties that SP have with attention^37–41^. We also note the possibility that what these findings indicate could potentially contribute to a disjointed and fragmented sense of reality, however a follow-up analysis did not indicate a significant correlation between “disorganization” and any of the meta-state statistics. Therefore it may be a more general relationship between a combination of positive symptoms that is related to the current results. Furthermore, it appears likely from our results here, combined with our previous work [34] that the way individuals travel through cognitive states may be considered on a continuum with HC toward one extreme, where transitions are smooth and occur at some optimum frequency, and SP spreading toward the other end of the spectrum as their positive symptoms increase, where transitions become frequent and abrupt (**Table 1 & Fig. 1**).

Our additional preliminary results, in which we find a trend in a data-driven k-means clustering based on meta-state statistics towards groupings reflective of symptomology, provide additional evidence that the metrics “Change between states” and “Total distance” are related to symptomology for SP. These two statistics appear to be the primary drivers of the clustering, as inferred from these statistics being the only ones which are significantly different between all 4 of the cluster groupings. This result is also of interest as potentially identifying a method of sensitively classifying individuals who have symptomology which overlaps disorder categories (e.g. SP and bipolar patients). However, we clearly need a larger sample size to verify these preliminary results.

Future work will investigate a larger group of patients, with a broader range of frequencies (i.e. including gamma band), and a longer scan duration. With a larger group of individuals we anticipate a definitive answer to the utility of our clustering approach based on meta-state statistics as a method of classifying patients. Higher frequencies and a longer scan duration may affect how many and what brain states are identified and how they vary over time. However, we would argue that scan duration should not dramatically affect the number of states, since only 3 states were identified in the current work. It is probable that the frequency of the neuronal oscillations will form different correlation patterns, which work together but are related in a complex manner. It is unknown at this time if this would affect the global meta-state level statistics, and this is an important open question. Lastly, we reiterate the importance of replicating these results with patients who have a broader range of symptoms, since there is no a priori reason to conclude that the results here will remain linear at the extreme.

